# Persistent Anhedonia After Intermittent Long-Access Nicotine Self-Administration in Rats

**DOI:** 10.1101/2025.03.19.644137

**Authors:** Ranjithkumar Chellian, Azin Behnood-Rod, Adriaan W. Bruijnzeel

## Abstract

Tobacco use disorder is a chronic condition characterized by compulsive nicotine use and withdrawal symptoms after smoking cessation. Smoking is the leading preventable cause of morbidity and mortality worldwide. Smoking cessation leads to anhedonia, which is an inability to experience pleasure from previously enjoyed activities and is caused by dysregulation of the brain’s reward and stress systems. It is also a key withdrawal symptom that contributes to relapse to smoking after a period of abstinence. To better understand the development of anhedonia, we investigated its onset and time course in rats that self-administered nicotine. Rats were implanted with intracranial self-stimulation (ICSS) electrodes to assess reward function and intravenous catheters for nicotine self-administration. Elevations in ICSS brain reward thresholds reflect decreased sensitivity to rewarding electrical stimuli, indicating anhedonia. The rats self-administered 0.06 mg/kg of nicotine intermittently, three days per week, for seven weeks. Brain reward thresholds were determined once a week 24 h after nicotine self-administration during weeks 1 to 3, and at 12, 24, and 48 h during weeks 4, 5, and 7. Elevations in brain reward thresholds were not observed during the first four weeks of nicotine self-administration. However, the brain reward thresholds were elevated in both weeks 5 and 7 at least 12 h after nicotine self-administration, indicating that anhedonia emerges gradually and then persists. As withdrawal severity gradually increases, smoking cessation may become more challenging. Therefore, behavioral or pharmacological interventions soon after smoking initiation are critical to prevent the development of a tobacco use disorder.

**Highlights:**

- Intermittent long-access nicotine self-administration leads to dependence in rats.
- Cessation of nicotine intake increased reward thresholds after 5 and 7 weeks.
- Anhedonia emerged after 5 weeks of long-access nicotine self-administration.
- Brain reward thresholds increased during spontaneous and precipitated withdrawal.
- Response latencies increased during both spontaneous and precipitated withdrawal.

## 1. Introduction

Tobacco use disorder is a chronic condition characterized by the compulsive use of tobacco products, withdrawal symptoms and cravings following smoking cessation, and relapse after periods of abstinence. Worldwide, there are about 1.25 billion people who smoke, and tobacco use is the number one preventable cause of morbidity and mortality (WHO, 2024). Tobacco use has been associated with a wide range of diseases, including cardiovascular disease, respiratory disease, cancer, and Alzheimer’s disease (Ambrose and Barua, 2004; Forey et al., 2011; Ott et al., 1998; Sasco et al., 2004). Furthermore, tobacco-related illnesses impose a significant economic burden on healthcare systems, costing hundreds of billions of dollars annually in medical expenses and lost productivity (Goodchild et al., 2018; Nargis et al., 2022). Despite extensive public health campaigns and the availability of FDA-approved smoking cessation aids, smoking and relapse rates remain high (Agboola et al., 2015; Cornelius et al., 2023; Williams et al., 2007).

Smoking has adverse health effects, and most smokers would like to quit, but fewer than 10% are able to do so each year (VanFrank, 2024). Smoking cessation leads to withdrawal symptoms that can hinder cessation efforts (Allen et al., 2008; McCarthy et al., 2006). Common withdrawal symptoms include nicotine cravings, irritability, anxiety, anhedonia, difficulty concentrating, increased appetite, and sleep disturbances (Hughes, 2007). These nicotine withdrawal symptoms typically begin a few hours after the last cigarette and peak within the first week of abstinence. Most people relapse to smoking during the first week of abstinence when withdrawal is most severe (Hughes et al., 2004; Jarvis, 2004). The FDA-approved smoking cessation drugs varenicline (Chantix), bupropion (Wellbutrin), and nicotine replacement therapies (NRTs) diminish nicotine withdrawal symptoms in humans and animal models of tobacco use (Chellian et al., 2021a; Rigotti et al., 2022). However, these treatments have adverse side effects and two thirds of people stop using them within the first three months (Cheng et al., 2024).

Understanding the neurobiological mechanisms underlying nicotine withdrawal is critical for developing effective behavioral and pharmacological smoking cessation strategies. Traditional studies have often used non-contingent nicotine administration methods, such as injections, osmotic minipumps, or passive exposure to tobacco smoke and nicotine vapor, to induce dependence and observe withdrawal symptoms (Chellian et al., 2021a). While these approaches have provided valuable insights, they do not fully capture the complexities of tobacco use disorder, as they do not induce the same neuronal adaptations as drug self-administration (Jacobs et al., 2003; Markou et al., 1999; Orrù et al., 2016; Steketee, 2017). Thus, animal models in which rats become dependent through nicotine self-administration have greater face and construct validity compared to models in which rats are non-contingently exposed to nicotine to induce dependence (Geyer and Markou, 1995; Willner, 1984).

Nicotine mediates its rewarding effects by activating α4β2, α6, α3β4, and α7 nicotinic acetylcholine receptors (nAChRs) in the brain, which in turn induces the release of dopamine and glutamate in the nucleus accumbens (Picciotto and Kenny, 2021). Nicotine withdrawal is associated with a decreased release of dopamine in the nucleus accumbens, as well as disrupted glutamate homeostasis (Li et al., 2014; Zhang et al., 2012). These nicotine-induced neuroadaptations in dopamine and glutamate signaling may contribute to anhedonia following cessation of nicotine intake. Furthermore, the activation of brain stress systems, including the release of corticotropin-releasing factor (CRF) and dynorphin, may further increase the anhedonia associated with nicotine withdrawal (Bruijnzeel, 2009; Bruijnzeel et al., 2007; Tejeda et al., 2012).

Very few studies have investigated the relationship between drug self-administration and brain reward function using the ICSS procedure. The ICSS procedure measures the sensitivity to rewarding electrical stimuli. The acute administration of drugs of abuse, such as nicotine, increases sensitivity to rewarding electrical stimuli and lowers brain reward thresholds (Igari et al., 2013). In contrast, cessation of long-term drug administration decreases sensitivity to rewarding electrical stimuli and increases brain reward thresholds (Epping-Jordan et al., 1998). Many studies have examined the effects of noncontingent drug exposure on ICSS thresholds (Chellian et al., 2021a), but few have assessed how drug self-administration affects brain reward function. Jang et al. reported that reward thresholds increased progressively with long access to methamphetamine, indicating a decrease in brain reward function (Jang et al., 2013). Furthermore, rats with long access to cocaine have elevated brain reward thresholds compared to rats with short access to cocaine (Ahmed et al., 2002). Harris et al. examined the effects of nicotine withdrawal on reward thresholds in rats following 22-h/day nicotine self-administration for approximately 6 weeks (∼43 sessions)(Harris et al., 2011). After this extended nicotine exposure period, reward thresholds were assessed during both antagonist-precipitated and spontaneous withdrawal. The study found that nicotine-dependent rats have elevated brain reward thresholds during spontaneous and precipitated withdrawal.

Previous studies have shown that cessation of noncontingent nicotine administration and cessation of long-term nicotine self-administration lead to anhedonia (Epping-Jordan et al., 1998; Harris et al., 2011). To study the role of anhedonia in nicotine intake, a better understanding of the time-dependent development of anhedonia is critical. Therefore, by using a long-access nicotine self-administration paradigm in combination with ICSS, we determined the time course of the development of anhedonia in animals self-administering nicotine. The rats were given intermittent access to nicotine as intermittent access has been shown to lead to higher levels of nicotine intake compared to daily access (Chellian et al., 2024; Cohen et al., 2012). The present findings show that rats that self-administer nicotine develop anhedonia after 4 weeks of intermittent long-access nicotine self-administration.

## 2. Material and methods

### 2.1. Animals

Adult male (200 - 250 g, 8-9 weeks of age; N=13) Wistar rats were purchased from Charles River (Raleigh, NC). The rats were housed in a climate-controlled vivarium on a reversed 12 h light-dark cycle (light off at 7 AM). The rats were singly housed after the implantation of the ICSS electrodes. Prior to the onset of the studies, food was available ad libitum in the home cage. During the food training and the nicotine self-administration sessions, the rats were fed 90-95 percent (23 g/day) of their ad libitum food intake. A mild level of food restriction facilitates food training and nicotine self-administration in rats (Donny et al., 1998; Garcia et al., 2014). Water was available ad libitum throughout the study. The experimental protocols were approved by the University of Florida Institutional Animal Care and Use Committee (IACUC). All experiments were performed in accordance with relevant IACUC guidelines and regulations and in compliance with ARRIVE guidelines 2.0 (Animal Research: Reporting of *In Vivo* Experiments).

### 2.2. Drugs

For intravenous self-administration, (-)-nicotine hydrogen tartrate (Sigma-Aldrich, St. Louis, MO, USA) was dissolved in sterile saline (0.9 % sodium chloride). The rats self-administered 0.03 or 0.06 mg/kg/inf of nicotine in a volume of 0.1 ml/inf. Nicotine doses are expressed as base. Mecamylamine hydrochloride (Sigma-Aldrich, St. Louis, MO, USA) was dissolved in sterile saline and administered subcutaneously (1 ml/kg). The mecamylamine doses are expressed as salt.

### 2.3. Experimental design

The rats were prepared with ICSS electrodes in the medial forebrain bundle and trained on the ICSS procedure. The rats were singly housed after the implantation of the ICSS electrodes. The rats were also trained to respond for food pellets (45 mg, F0299, Bio-Serv, Frenchtown, NJ) in operant self-administration chambers under a fixed ratio 1 (FR1) and an FR1 time-out 10 s (FR1-TO10) schedule. Food training was done in the morning, and ICSS training in the afternoon. The rats were fed (23 g) immediately after the food training sessions. The food training sessions were conducted for 10 days. After completing the last food training sessions, the rats were fed ad libitum and ICSS training continued until the brain reward thresholds stabilized. After completing ICSS training, the rats were prepared with catheters in the jugular vein. Baseline brain reward thresholds and response latencies were determined on three consecutive days before the implantation of the catheters. One day before the start of the nicotine self-administration sessions, the rats were allowed to respond for food pellets under the FR1-TO10s schedule (one 20-min session), followed by an ICSS session.

The rats were allowed to self-administer nicotine for five daily 1 h baseline sessions. During the first three sessions (days 1-3), the rats self-administered 0.03 mg/kg/inf of nicotine under an FR1-TO10s schedule. During the following two sessions (days 4 and 5) the rats self-administered 0.06 mg/kg/inf of nicotine under an FR1-TO60s schedule. The 0.06 mg/kg/inf dose of nicotine leads to higher levels of nicotine intake than the 0.03 mg/kg/inf dose (Chaudhri et al., 2005; Shoaib and Stolerman, 1999). During the first day that the rats received the 0.03 or 0.06 mg/kg/inf dose, nicotine intake was limited to prevent aversive effects (i.e., seizures and negative mood state). The maximum number of infusions was set to 20 on the first day that the rats received the 0.03 mg/kg/inf dose and to 10 on the first day that the rats received the 0.06 mg/kg/inf dose. After five daily nicotine self-administration sessions, the rats were allowed to self-administer nicotine (0.06 mg/kg/inf) under an intermittent long-access schedule (23 h/day). The rats self-administered nicotine during the dark (11 h) and the light (12 h) phase. During the light phase, the cubicle was illuminated by a house light (ENV-229M, Med Associates, St. Albans, VT) that was mounted on the ceiling of the cubicle. The light was programmed to turn on automatically and remained on for the entire duration of the light phase. The light intensity in the operant chambers during the light phase was 13 lux. Intermittent long-access nicotine self-administration sessions were conducted three days per week (Monday, Wednesday, and Friday) for seven weeks (21 sessions). Responding on the active lever resulted in the delivery of a nicotine infusion (0.1 ml infused over a 6.5-s period). The initiation of the delivery of an infusion was paired with a cue light, which remained illuminated throughout the time-out period. Responding on the inactive lever was recorded but did not have scheduled consequences. The active and inactive lever were retracted during the time-out period. During the self-administration period, the rats received 90-95 percent (23 g) of their normal ad libitum food intake in the home cage immediately after the operant sessions. In addition, the rats received water and food (23 g) in the operant chamber on long-access self-administration days.

During the seven-week long-access nicotine self-administration period, the effects of spontaneous nicotine withdrawal and mecamylamine-precipitated nicotine withdrawal on brain reward thresholds and response latencies in the ICSS procedure were investigated. Spontaneous withdrawal was assessed after the 3rd (week 1), 6th (week 2), 9th (week 3), 12th (week 4), 15th (week 5), and 21st (week 7) long-access nicotine self-administration sessions. In weeks 1 – 3, the ICSS sessions were conducted immediately after (0 h) and 24 h after the long-access nicotine self-administration sessions. In weeks 4, 5, and 7, the ICSS sessions were conducted immediately after (0 h), 12 h, 24 h, and 48 h after the long-access nicotine self-administration sessions. Mecamylamine-precipitated withdrawal was assessed immediately after the 16th - 19th (weeks 6 and 7) long-access nicotine self-administration sessions. Mecamylamine (0, 0.5, 1.5, and 3 mg/kg, SC) was administered according to a Latin square design. Mecamylamine was administered 15 min before the ICSS sessions. The mecamylamine dose was based on previous studies in which we investigated the effects of mecamylamine on reward thresholds in nicotine exposed rats (Bruijnzeel et al., 2010; Geste et al., 2020).

### 2.4. Electrode implantation and ICSS procedure

The rats were prepared with ICSS electrodes in the medial forebrain bundle and trained on the ICSS procedure (Chellian et al., 2021b; Tan et al., 2019; Xue et al., 2018). Briefly, the rats were anesthetized with an isoflurane and oxygen vapor mixture (1-3%) and placed in a stereotaxic frame (David Kopf Instruments, Tujunga, CA, USA). Electrodes were implanted in the medial forebrain bundle with the incisor bar set 5 mm above the interaural line (anterior-posterior -0.5 mm, medial-lateral ±1.7 mm, dorsal-ventral -8.3 mm from dura). The rats were trained on a modified discrete-trial ICSS procedure in operant conditioning chambers (Med Associates, Georgia, VT, USA), housed within sound-attenuating enclosures. A five cm wide metal response wheel was centered on a sidewall, with a photobeam detector recording every 90 degrees of rotation. Brain stimulation was delivered by constant current stimulators (Model 1200C, Stimtek, Acton, MA, USA). The rats were trained to turn the wheel on a fixed ratio 1 (FR1) schedule of reinforcement. Each quarter-turn of the wheel resulted in the delivery of a 0.5-second train of 0.1 ms cathodal square-wave pulses at a frequency of 100 Hz. After acquiring responding for stimulation on the FR1 schedule (100 reinforcements within 10 minutes), the rats were trained on a discrete-trial current-threshold procedure. The discrete-trial current-threshold procedure is a modification of a task developed by Kornetsky and Esposito (Kornetsky and Esposito, 1979), and previously described in detail (Chellian et al., 2021b; Tan et al., 2019; Xue et al., 2018). Each trial began with the delivery of a noncontingent electrical stimulus, followed by a 7.5-s response window during which the animals could respond for a second identical stimulus. A response during this 7.5-s window was labeled a positive response, and the lack of a response was labeled a negative response. During the 2-s period immediately after a positive response, additional responses had no consequences. The inter-trial interval (ITI), which followed a positive response or the end of the response window, had an average duration of 10 s (ranging from 7.5 to 12.5 s). Responses during the ITI resulted in a 12.5-s delay of the onset of the next trial. During the training sessions, the duration of the ITI and delay periods induced by time-out responses were increased until the animals performed consistently at standard test parameters. The training was completed when the animals responded correctly to more than 90% of the noncontingent electrical stimuli. It took 2-3 weeks of training for most rats to meet this response criterion. The rats were then tested on the current-threshold procedure in which stimulation intensities varied according to the classical psychophysical method of limits. Each test session consisted of four alternating series of descending and ascending current intensities starting with a descending sequence. Blocks of three trials were presented to the rats at a given stimulation intensity, and the intensity was altered between blocks of trials by 5 µA steps. The initial stimulus intensity was set 30 µA above the baseline current threshold for each animal. Each test session typically lasted 30-40 min and provided two dependent variables for behavioral assessment (brain reward thresholds and response latencies). The brain reward threshold (µA) was defined as the midpoint between stimulation intensities that supported responding and stimulation intensities that failed to support responding. The response latency was defined as the time interval between the beginning of the noncontingent stimulus and a response. A decrease in reward thresholds is indicative of enhanced reward function (Kornetsky and Esposito, 1979). Drugs with sedative effects increase the response latencies, and stimulants decrease the response latencies (Igari et al., 2013; Liebman, 1985).

### 2.5. Food training

Rats were trained to press a lever for food pellets in operant chambers that were placed in sound- and light-attenuated cubicles (Med Associates, St. Albans, VT). Responding on the active lever resulted in the delivery of a food pellet (45 mg, F0299, Bio-Serv, Frenchtown, NJ), and responding on the inactive lever was recorded but did not have scheduled consequences. Food delivery was paired with a cue light, which remained illuminated throughout the time-out period. The food training sessions were conducted for 10 days. Instrumental training started under a FR1-TO1s reinforcement schedule, and the rats remained on this schedule for 5 days (30 min sessions per day). On day 6, the time-out period was increased to 10 s. The rats were allowed to respond for food pellets under the FR1-TO10s schedule (20 min sessions) for 5 days. Both levers were retracted during the 10 s time-out period.

### 2.6. Intravenous catheter implantation

The catheters were implanted as described before (Chellian et al., 2021b; Chellian et al., 2021c; Chellian et al., 2022). The rats were anesthetized with an isoflurane-oxygen vapor mixture (1-3%) and prepared with a catheter in the right jugular vein. The catheters consisted of polyurethane tubing (length 10 cm, inner diameter 0.64 mm, outer diameter 1.0 mm, model 3Fr, Instech Laboratories, Plymouth Meeting, PA). The right jugular vein was isolated, and the catheter was inserted 2.9 cm. The tubing was then tunneled subcutaneously and connected to a vascular access button (Instech Laboratories, Plymouth Meeting, PA). The button was exteriorized through a 1-cm incision between the scapulae. During the 7-day recovery period, the rats received daily infusions of the antibiotic Gentamycin (4 mg/kg, IV, Sigma-Aldrich, St. Louis, MO). A sterile heparin solution (0.1 ml, 50 U/ml) was flushed through the catheter before and after administering the antibiotic and after nicotine self-administration. After flushing the catheter, 0.05 ml of a sterile heparin/glycerol lock solution (500 U/ml, Instech Laboratories, Plymouth Meeting, PA) was infused into the catheter. The animals received carprofen (5 mg/kg, SC) daily for 72 h after the surgery. Catheter patency was evaluated by infusing 0.2 ml of the ultra-short-acting barbiturate Brevital (1% methohexital sodium). Rats with patent catheters displayed a sudden loss of muscle tone. If the rats did not respond to Brevital, their self-administration data were excluded from the analysis. Three rats did not respond to Brevital during the third and fourth weeks of long-access nicotine self-administration, and one rat did not respond to Brevital at the end of the study. Therefore, data from four rats were excluded.

### 2.7. Statistics

One-way ANOVAs were used to compare absolute brain reward thresholds and response latencies after catheter surgery to the 3-day baseline average recorded before surgery. The brain reward thresholds and the response latencies during nicotine withdrawal were expressed as a percentage of the 3-day baseline that was obtained prior to the catheter surgery. The percentage change in thresholds and latencies were analyzed with one-way ANOVAs, with withdrawal as within-subjects factor. Baseline (1 h/day, 5 days) and long-access (23 h/day, 21 sessions) nicotine self-administration data were analyzed using one- or two-way ANOVAs with lever (active versus inactive) and session as within-subject factors. For all statistical analyses, significant interaction effects found in the ANOVAs were followed by Bonferroni’s post hoc tests to determine which groups differed. P-values less than or equal to 0.05 were considered significant. Data were analyzed with IBM SPSS Statistics version 29 and GraphPad Prism version 10.1.2. The figures were generated using GraphPad Prism version 10.1.2.

## 3. Results

### 3.1. Baseline nicotine self-administration

The rats self-administered 0.03 mg/kg/inf of nicotine for three days (1 h/day) and 0.06 mg/kg/inf of nicotine for 2 days (1 h/day). During the first three nicotine self-administration sessions (0.03 mg/kg/inf), the rats responded more on the active lever than on the inactive lever (Fig S1A; Lever F1,8=30.447, P < 0.001). Active and inactive lever responses did not change over time (Fig S1A; Session F2,16=1.079, NS; Lever × Session F2,16=3.438, NS). Nicotine intake slightly decreased across these sessions (Fig S1B; Session F2,16=3.806, P < 0.05). During the following two self-administration sessions (0.06 mg/kg/inf), active lever responses remained higher than inactive lever responses and increased over time, whereas inactive lever responses decreased over time (Fig S1C; Lever F1,8=43.859, P < 0.001; Session F1,8=13.369, P < 0.01; Lever × Session F1,8=9.166, P < 0.05). Nicotine intake also increased during these sessions (Fig S1D; Session F1,8=12.815, P < 0.01).

### 3.2. Intermittent long-access nicotine self-administration

After the baseline sessions, the rats self-administered nicotine under an intermittent long-access schedule for 21 sessions. During the 21 sessions (0.06 mg/kg/inf, 23 h/session, 3 sessions/week), the rats responded more on the active lever than on the inactive lever (Fig 1A; Lever F1,8=51.623, P < 0.001). Responding on the active lever decreased over time, while the responding on the inactive lever initially increased and then decreased (Fig 1A; Session F20,160=4.071, P < 0.001; Lever × Session F20,160=2.275, P < 0.01). Nicotine intake decreased over the 21 sessions (Fig 1B; Session F20,160=5.719, P < 0.001).

**Figure 1.**
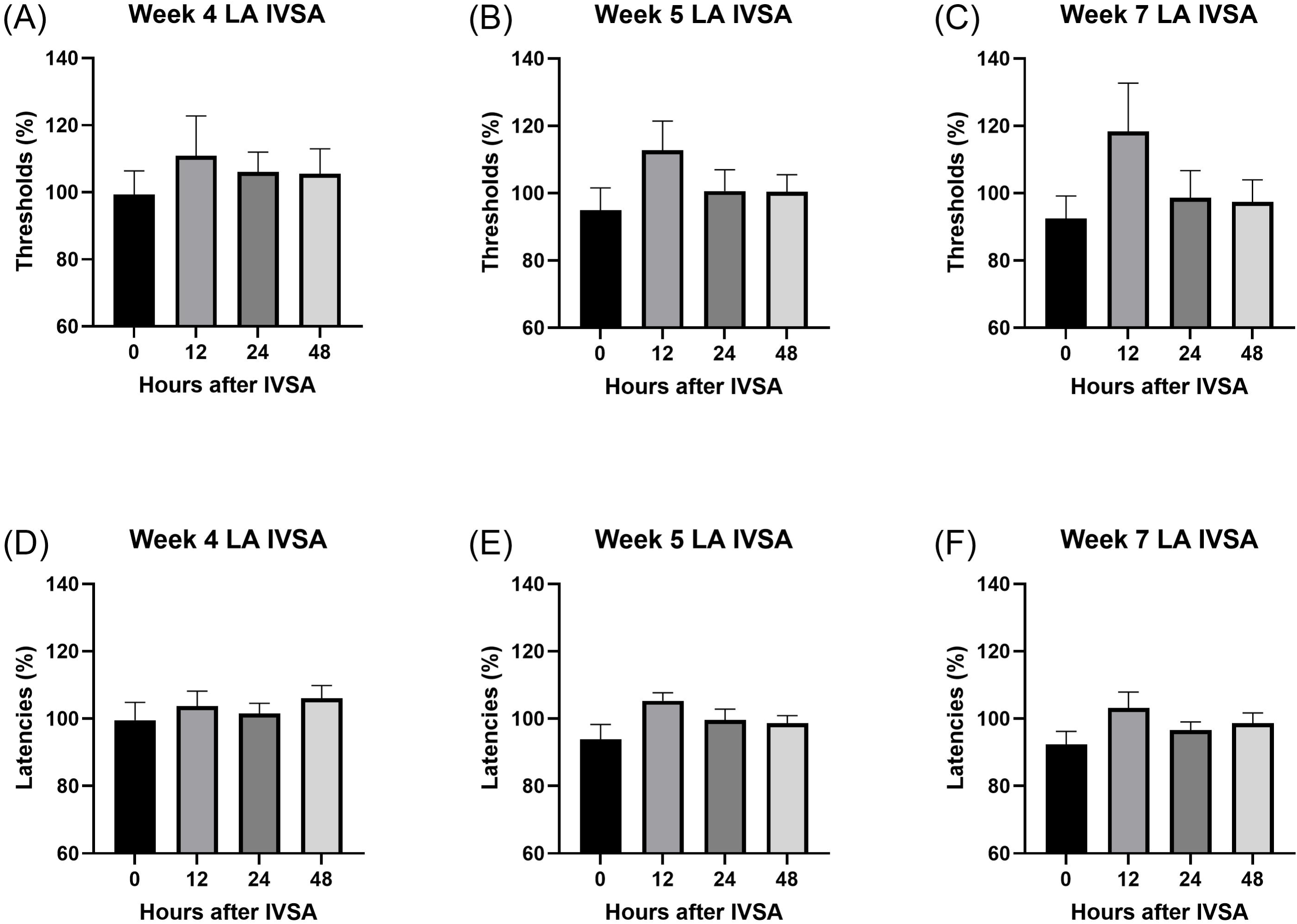
Nicotine self-administration in male rats under an intermittent long-access self-administration paradigm. The rats self-administered nicotine (0.06 mg/kg/inf) in 23-h self-administration sessions for 21 sessions. (A) Active and inactive lever responses during nicotine self-administration. Rats responded significantly more on the active lever than on the inactive lever across sessions. (B) Nicotine intake and the number of nicotine infusions across sessions. N = 9/group. Data are expressed as means ± SEM.

### 3.3. Spontaneous nicotine withdrawal and the brain reward function

#### Reward thresholds

Before the start of nicotine self-administration, there was no difference in the absolute brain reward thresholds compared to the 3-day baseline reward thresholds (Fig S2A; F1,8=0.046, NS). During the first four weeks of long-access nicotine self-administration, nicotine withdrawal did not affect the brain reward thresholds (Fig S3A, week 1, F1,8=0.066, NS; Fig S3B, week 2, F1,8=0.703, NS; Fig S3C, week 3,F1,8=3.304, NS; Fig 2A, week 4, F3,24=0.591, NS). However, in weeks 5 and 7, nicotine withdrawal elevated the brain reward thresholds (Fig 2B, week 5, F3,24=3.785, P < 0.05; Fig 2C,week 7, F3,24=3.464, P < 0.05).

**Figure 2.**
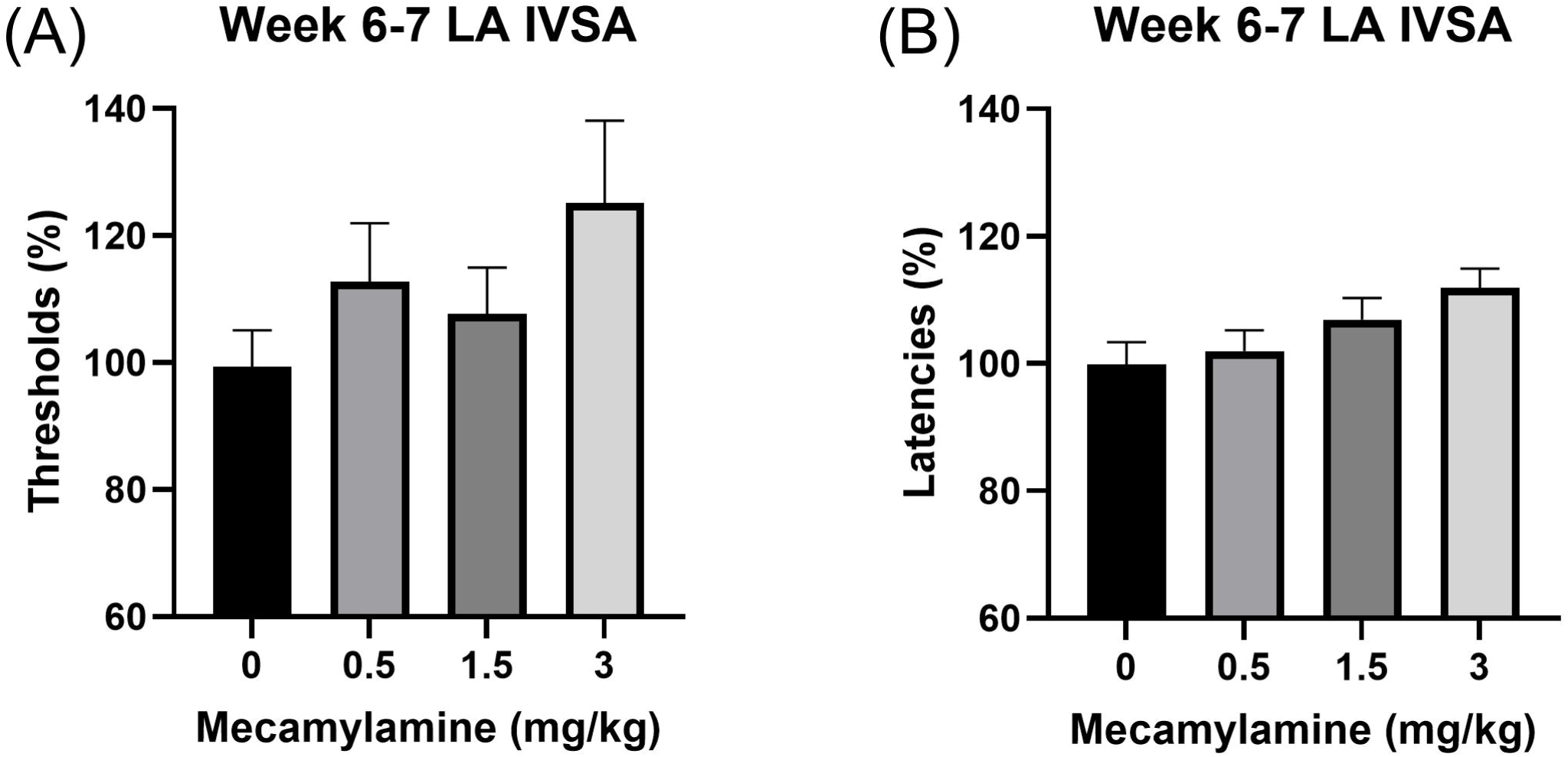
Effects of nicotine withdrawal on brain reward thresholds and response latencies following long-access intravenous self-administration. The rats self-administered nicotine (0.06 mg/kg/inf) in 23-h self-administration sessions. (A–C) Brain reward thresholds were assessed at 0, 12, 24, and 48 h after nicotine self-administration during weeks 4, 5, and 7. (A) Week 4: No significant changes in brain reward thresholds were observed. (B) Week 5 and (C) Week 7: Cessation of nicotine intake significantly elevated brain reward thresholds. (D–F) Response latencies were assessed at 0, 12, 24, and 48 h after nicotine self-administration during weeks 4, 5, and 7. (D) Week 4: No significant changes in response latencies were observed. (E) Week 5 and (F) Week 7: Cessation of nicotine intake significantly increased response latencies. N = 9/group. Data are expressed as means ± SEM.

#### Response latencies

Before the onset of nicotine self-administration, there was no difference in the absolute response latencies compared to the 3-day baseline response latencies (Fig S2B; F1,8=0.273, NS). During the first four weeks of long-access nicotine self-administration, cessation of nicotine intake did not affect the response latencies (Fig S3D, week 1, F1,8=0.13, NS; Fig S3E, week 2, F1,8=1.142, NS; Fig S3F, week 3, F1,8=0.991, NS; Fig 2D, week 4, F3,24=0.653, NS). However, in weeks 5 and 7, cessation of nicotine intake increased the response latencies (Fig 2E, week 5, F3,24=3.813, P < 0.05; Fig 2F,week 7, F3,24=3.157, P < 0.05).

### 3.4. Mecamylamine-precipitated nicotine withdrawal and the brain reward function

#### Reward thresholds and response latencies

After long-access nicotine self-administration, treatment with mecamylamine elevated both the brain reward thresholds (Fig 3A; F3,24=4.342, P < 0.05) and increased the response latencies (Fig 3B; F3,24=8.372, P < 0.001).

**Figure 3.**
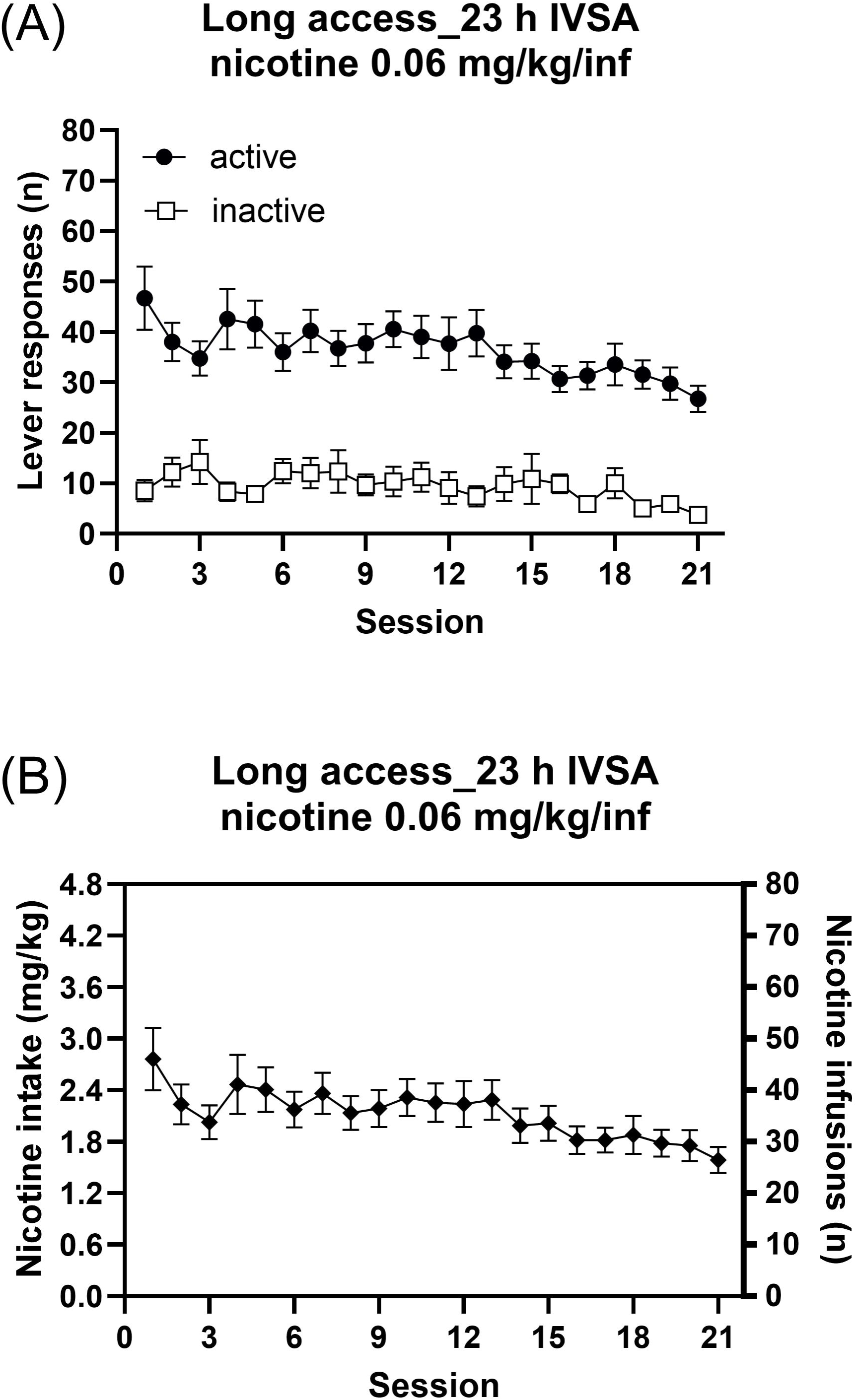
Mecamylamine-precipitated withdrawal increase brain reward thresholds and response latencies in rats following long-access nicotine self-administration. The rats self-administered nicotine (0.06 mg/kg/inf) in 23-h intravenous self-administration sessions for 6–7 weeks. (A) Brain reward thresholds following mecamylamine administration. Mecamylamine significantly increased brain reward thresholds. (B) Response latencies following mecamylamine administration. Mecamylamine significantly increased response latencies. N = 9/group. Data are expressed as means ± SEM.

## 4. Discussion

The goal of the present study was to investigate the time course of the development of anhedonia during abstinence in rats that self-administer nicotine. The rats self-administered nicotine 3 times per week in 23-h session and withdrawal was investigated between weeks 1 and 7. During the first four weeks of long-access nicotine self-administration, cessation of nicotine intake did not alter the brain reward thresholds. However, in weeks 5 and 7, cessation of nicotine intake led to significant elevations in brain reward thresholds. This indicates that the rats develop anhedonia after the cessation of nicotine self-administration. Furthermore, the administration of the nAChR antagonist mecamylamine elevated the brain reward thresholds of the rats following long-access nicotine self-administration. Taken together, these findings indicate that animals that self-administer nicotine slowly develop nicotine dependence over time. This work has clinical implications and suggests that early interventions in new smokers and vapers may mitigate the development of withdrawal-induced anhedonia and reduce the risk of relapse.

In the present study, we investigated the development of dependence in animals that self-administered 0.06 mg/kg/inf of nicotine. This dose is somewhat higher than the standard 0.03 mg/kg/inf dose, which is commonly used in rat nicotine self-administration studies (Bruijnzeel and Markou, 2003; Corrigall and Coen, 1989). However, an advantage of this higher dose is that it leads to greater nicotine intake compared to rats self-administering lower doses of nicotine and may hasten the development of dependence. Animals with short access to nicotine self-administer more nicotine when given access to 0.06 mg/kg/inf compared to 0.01 or 0.03 mg/kg/inf of nicotine (Chaudhri et al., 2005; Shoaib and Stolerman, 1999). On a similar note, O’Dell et al. showed that rats self-administer more nicotine in a long-access paradigm when they have access to higher compared to lower doses of nicotine (O’Dell et al., 2007). Furthermore, in one of our previous studies, rats with intermittent long access to 0.03 mg/kg/inf of nicotine had an intake of approximately 1.4 mg/kg of nicotine (Geste et al., 2020). In contrast, in the present study, rats self-administered about 2.4 mg/kg of nicotine, indicating that nicotine intake was approximately 40% higher when the animals had access to 0.06 versus 0.03 mg/kg of nicotine.

The present findings demonstrate that prolonged nicotine self-administration leads to a delayed increase in brain reward thresholds, which was being observed after 5 and 7 weeks of nicotine self-administration. This pattern aligns with prior research on other drugs of abuse, such as cocaine, where long-access self-administration progressively elevates ICSS thresholds (Ahmed et al., 2002). Similarly, long-access methamphetamine self-administration has been reported to cause elevations in brain reward thresholds (Jang et al., 2013). However, one major difference between our nicotine study and the studies with cocaine and methamphetamine was the number of self-administration sessions required before elevations in brain reward thresholds were observed. In the present nicotine study, elevations in brain reward thresholds were observed after twelve nicotine self-administration sessions, whereas in the cocaine study, elevations were observed after just four sessions, and similarly, elevations were seen after four methamphetamine self-administration sessions. There are several possible explanations for the difference in the time course of the development of anhedonia between rats self-administering nicotine versus cocaine and methamphetamine. One possibility is that differences in the potency and pharmacokinetics of nicotine compared to cocaine and methamphetamine contribute to the slower onset of anhedonia. Nicotine induces a smaller increase in dopamine and is not as rewarding as cocaine and methamphetamine (Benwell and Balfour, 1992; Igari et al., 2013; Izawa et al., 2006; Kenny et al., 2003; Radke et al., 2016; Sambo et al., 2017). Therefore, the neuroadaptations associated with nicotine exposure that lead to anhedonia might develop more gradually than those associated with exposure to cocaine and methamphetamine.

The outcome of our study aligns with the findings of Harris et al., who investigated anhedonia with the ICSS procedure in male rats following an extended period of nicotine self-administration (Harris et al., 2011). Specifically, they examined the effects of daily extinction sessions on ICSS thresholds in rats that self-administered 0.06 mg/kg/infusion of nicotine for about 43 sessions under a 22-h/day access schedule. They observed significant elevations in ICSS thresholds during the first three extinction sessions compared to nicotine-naïve controls that did not undergo nicotine self-administration. Our work extends upon their findings by showing that rats develop anhedonia following a much shorter self-administration period. Furthermore, in the study by Harris et al., the animals were tested in the ICSS procedure daily after an extinction session, whereas in our study, the rats were not placed in the self-administration chambers before ICSS testing and were instead tested at 12, 24, and 48-h following nicotine cessation (Harris et al., 2011). Interestingly, we found the most severe withdrawal effects at the 12-h time point, while Harris et al. did not assess brain reward thresholds at this early time point. Therefore, it is possible that they may have observed larger elevations in brain reward thresholds had they tested at earlier time points. This finding highlights the importance of assessing withdrawal at multiple time points to capture the time-dependent progression of anhedonia following nicotine cessation.

In the present study, we also investigated the effects of mecamylamine on brain reward thresholds and response latencies in rats that self-administered nicotine. The doses of mecamylamine used in this study do not affect brain reward thresholds or response latencies in rats that have not been exposed to nicotine (Bruijnzeel et al., 2009; Small et al., 2010). In the present study, we observed that mecamylamine elevated brain reward thresholds. This pattern of results suggests that long-access nicotine intake induces adaptations in nAChR signaling that contribute to the anhedonia associated with nicotine withdrawal (Besson et al., 2007). Mecamylamine also increased response latencies. Although response latencies are increased following the administration of sedative drugs (Liebman, 1985), these increases are also observed during drug withdrawal, including nicotine withdrawal (Bruijnzeel et al., 2009; Small et al., 2010; Spielewoy and Markou, 2003). In the case of nicotine withdrawal, the increase in response latencies may reflect a decreased motivation to obtain the rewarding electrical stimuli.

In conclusion, this study provides important insights into the time-dependent development of anhedonia in rats with intermittent long access to nicotine. The findings demonstrate that intermittent long-access nicotine self-administration leads to dependence, as indicated by spontaneous withdrawal-induced elevations in brain reward thresholds. The fact that mecamylamine precipitated withdrawal supports the idea that chronic nicotine self-administration induces adaptations in nAChR function which then contributes to withdrawal-induced anhedonia. This study highlights the delayed development of nicotine dependence and withdrawal in animals that self-administer nicotine. The extended time-period before dependence manifests in these animals provides an opportunity to investigate the neurobiological mechanisms that drive nicotine intake before and after the development of dependence. Based on prior studies with dependent animals, it could be hypothesized that stress systems play a greater role in nicotine intake after the development of dependence than before the development of dependence (Bruijnzeel et al., 2007; Cohen and George, 2013). These findings suggest that early interventions aimed at reducing smoking and vaping could help prevent the development of dependence and withdrawal-induced anhedonia, thereby increasing the likelihood of a successful smoking cessation attempt.

## Declaration of generative AI and AI-assisted technologies in the writing process

The authors used ChatGPT-4o for spelling and grammar checks. The authors wrote, reviewed, and edited the manuscript and take full responsibility for its final content.

## CRediT authorship contribution statement

**R. Chellian:** Formal analysis, Investigation, Writing - Review & Editing, Visualization. **A. Behnood-Rod:** Investigation, Project administration. **A. Bruijnzeel:** Conceptualization, Formal analysis, Writing - Original Draft, Visualization, Supervision, Project administration, Funding acquisition.

