## Supplemental figures S1-3 for "Persistent Anhedonia After Intermittent Long-Access Nicotine Self-Administration in Rats"

**Figure S1**

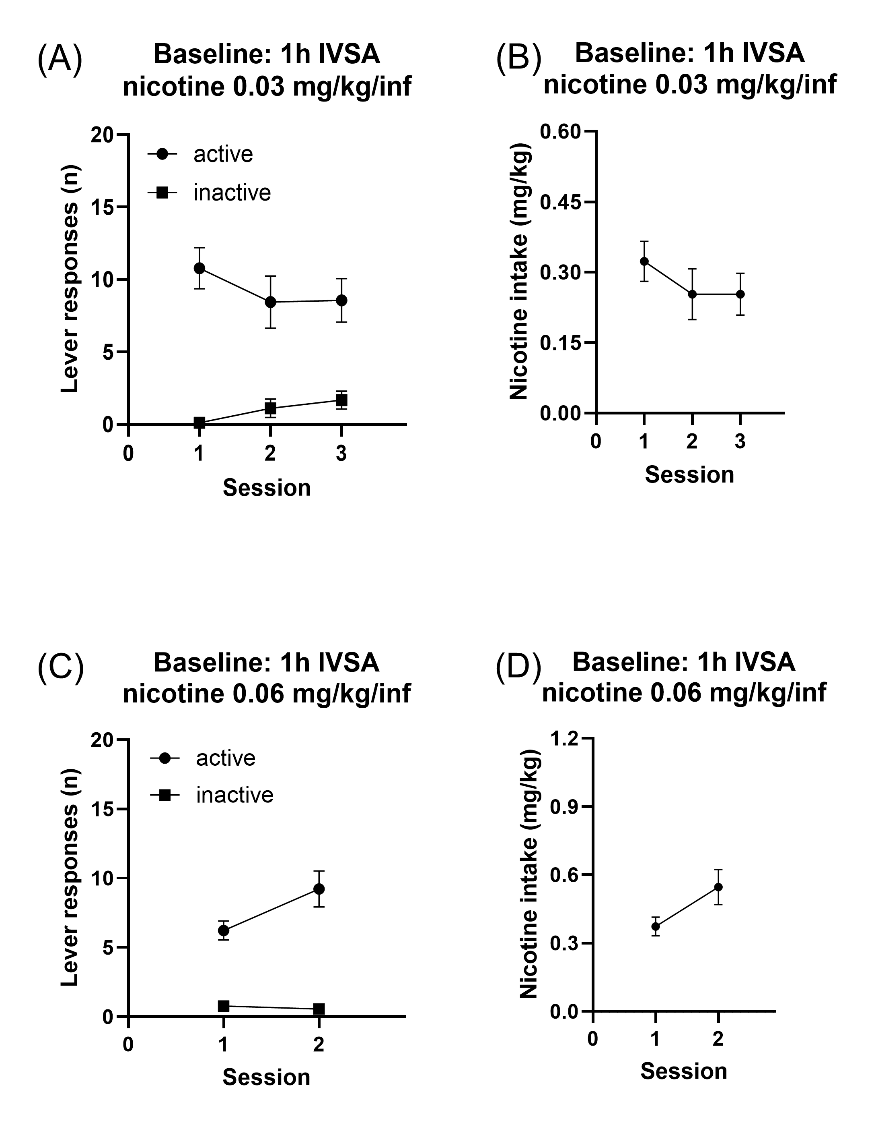

**Figure S1. Baseline nicotine self-administration in male rats during 1-hour intravenous self-administration (IVSA) sessions.** The rats self-administered nicotine at 0.03 mg/kg/inf (A, B) or 0.06 mg/kg/inf (C, D) in 1-hour sessions. The figures (A, C) show the number of active and inactive lever presses per session. Rats responded significantly more on the active lever than the inactive lever for both doses (A: F1,8 = 30.447, P < 0.001; C: F1,8 = 43.859, P < 0.001). Nicotine intake (B, D) slightly decreased across sessions for 0.03 mg/kg/inf (B: F2,16 = 3.806, P < 0.05) and increased for 0.06 mg/kg/inf (D: F1,8 = 12.815, P < 0.01). N = 9/group. Data are expressed as means ± SEM.

**Figure S2**

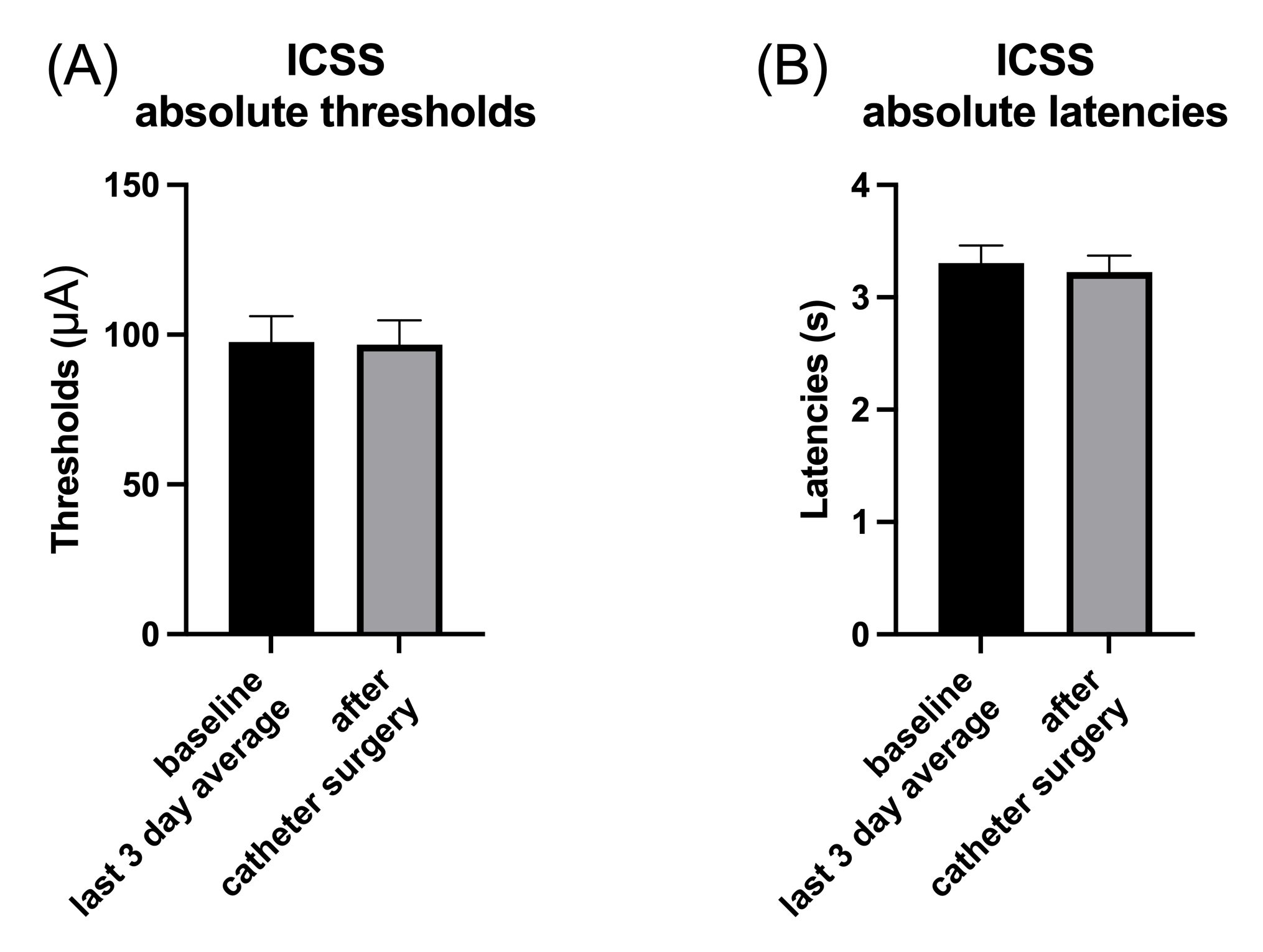

**Figure S2. Catheter surgery does not affect brain reward thresholds and response latencies.** The rats underwent ICSS testing before and after catheter implantation surgery. Absolute brain reward thresholds (A) and response latencies (B) were measured before the catheter surgery and after recovery from the surgery. There were no significant differences in brain reward thresholds before and after the surgery (F1,8 = 0.046, NS). There were also no differences between latencies before and after the surgery (F1,8 = 0.273, NS). These results indicate that catheter implantation did not significantly affect baseline ICSS performance. N = 9/group. Data are expressed as means ± SEM.

**Figure S3**

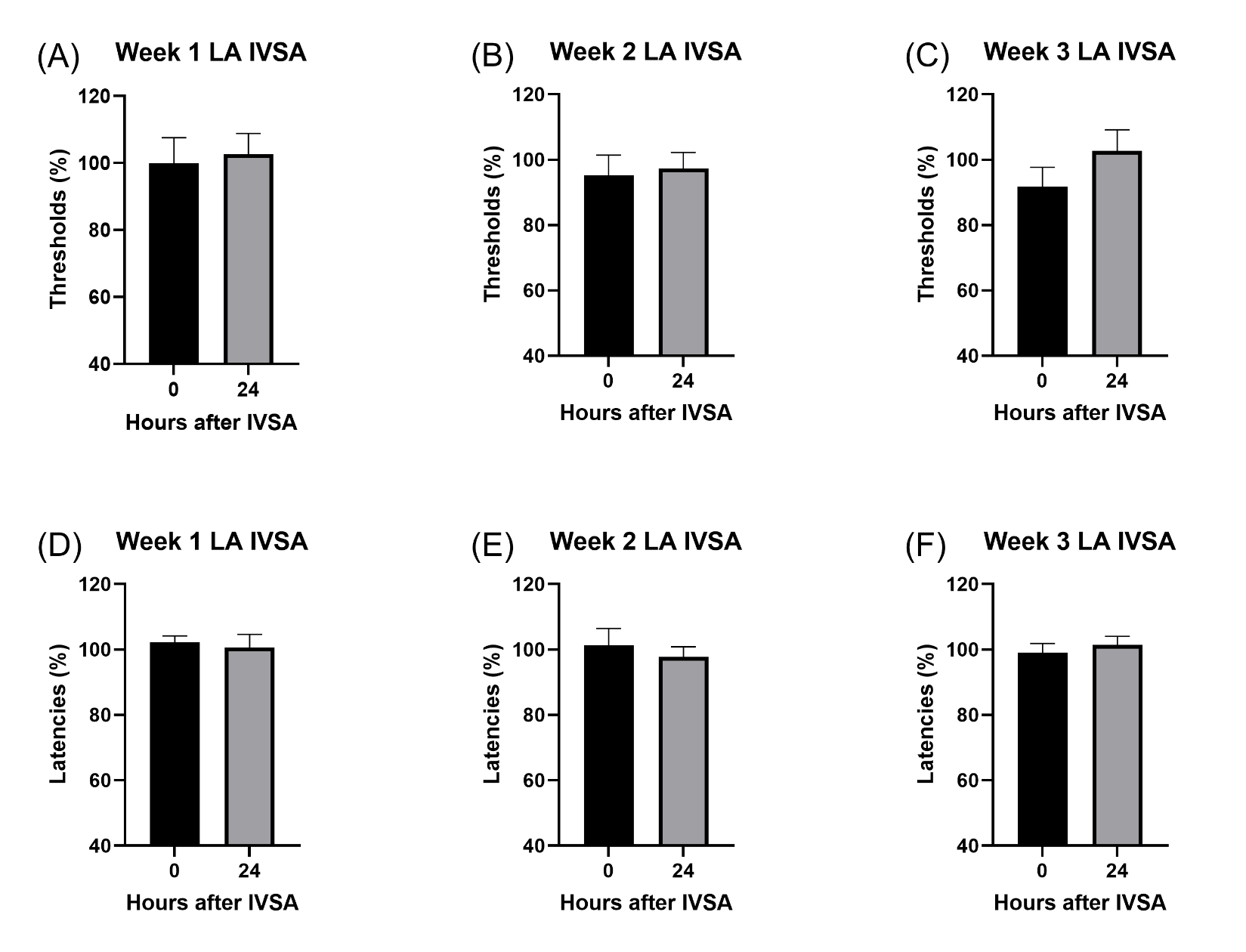

**Figure S3. Brain reward thresholds and response latencies during the first three weeks of long-access intravenous nicotine self-administration.** The rats self-administered nicotine (0.06 mg/kg/inf) in 23-hour self-administration sessions. Brain reward thresholds (A-C) were measured at 0 and 24 hours after nicotine self-administration in week 1 (A), week 2 (B), and week 3 (C). No significant effects of withdrawal on brain reward thresholds were observed (F1,8 = 0.066, NS; F1,8 = 0.703, NS; F1,8 = 3.304, NS, respectively). Response latencies (D-F) were also measured at 0 and 24 hours after nicotine self-administration in week 1 (D), week 2 (E), and week 3 (F), with no significant effects observed (F1,8 = 0.13, NS; F1,8 = 1.142, NS; F1,8 = 0.991, NS, respectively). These findings indicate that withdrawal-induced anhedonia did not develop during the first three weeks of nicotine self-administration. LA, long-access. N = 9/group. Data are expressed as means ± SEM.
